# High Throughput Variant Libraries and Machine Learning Yield Design Rules for Retron Gene Editors

**DOI:** 10.1101/2024.07.08.602561

**Authors:** Kate D. Crawford, Asim G. Khan, Santiago C. Lopez, Hani Goodarzi, Seth L. Shipman

## Abstract

The bacterial retron reverse transcriptase system has served as an intracellular factory for single-stranded DNA in many biotechnological applications. In these technologies, a natural retron non-coding RNA (ncRNA) is modified to encode a template for the production of custom DNA sequences by reverse transcription. The efficiency of reverse transcription is a major limiting step for retron technologies, but we lack systematic knowledge of how to improve or maintain reverse transcription efficiency while changing the retron sequence for custom DNA production. Here, we test thousands of different modifications to the retron-Eco1 ncRNA and measure DNA production in pooled variant library experiments, identifying regions of the ncRNA that are tolerant and intolerant to modification. We apply this new information to a specific application: the use of the retron to produce a precise genome editing donor in combination with a CRISPR-Cas9 RNA-guided nuclease (an editron). We use high-throughput libraries in *S. cerevisiae* to additionally define design rules for editrons. We extend our new knowledge of retron DNA production and editron design rules to human genome editing to achieve the highest efficiency retron-Eco1 editrons to date.

## INTRODUCTION

Retron components are increasingly being exploited for biotechnology due to their ability to produce DNA on demand in cells. In bacteria, retrons are a tripartite anti-phage system composed of a reverse transcriptase (RT), a non-coding RNA (ncRNA) that is reverse transcribed into DNA (RT-DNA), and an effector protein^1,2^. In biotechnology, the retron RT is used to reverse transcribe modified forms of retron ncRNA into RT-DNA that has been used as: donor DNA for precise editing in bacteria^3–8^, bacteriophage^9–11^, plants^12,13^, and eukaryotes^5,14–17^; DNA barcodes to record molecular events^18,19^; DNA containing transcription factor motifs for transcription factor activity attenuation^20^; DNA aptamers^21^; and DNAzymes for mRNA cleavage^22^.

Previous work has demonstrated that the abundance of retron reverse-transcribed DNA directly impacts the efficiency of downstream biotechnological applications. Specifically, modifications to the retron that generate more RT-DNA increase the efficiency of precise editing and the efficiency of event recording into a molecular ledger^15,18,20^. These previous works used the same modification to the retron ncRNA for increased RT-DNA production – extension of the a1/a2 region. However, the retron ncRNA has not been systematically interrogated to determine which elements are necessary, which are tolerant to modifications, and where it may be possible to increase reverse transcription beyond the endogenous element.

In the context of precise genome editing technologies, there are additional parameters that have not been investigated systematically. An editron, which combines retron components with CRISPR-Cas9 components to generate both a programmed double-strand break and the reverse transcribed donor to precisely repair it, has many degrees of freedom. These include among others, how to arrange the donor and gRNA relative to each other, where to situate the edit within the donor, or how long of a donor to use. Without a set of clear design rules, users are left to either empirically test many designs for their desired edit or pick an arbitrary design which may not perform optimally.

To rectify this lack of systematic investigation, we comprehensively tested all parameters of the retron ncRNA for their effect on RT-DNA production in high throughput, used these findings to build a machine learning model of RT-DNA production, and used the output of the model to inform high-throughput tests of editing parameters in yeast. Finally, we extended these findings to human cells, resulting in a set of design rules for RT-DNA production and retron-based editing that apply broadly.

## MATERIALS AND METHODS

Biological replicates were taken from distinct samples, not the same sample measured repeatedly. For *E. coli* variant libraries, each biological replicate is an independent electroporation and expression of the libraries into the strain bSLS.114. For *S. cerevisiae* variant libraries, each biological replicate is an independent transformation and expression of the variant libraries using a scaled-up version of the Zymo Frozen-EZ Yeast Transformation II Kit into the respective yeast strains containing the editing site. For human validation, each biological replicate is an independent transfection and expression of variants using Lipofectamine 3000 into a Cas9-containing HEK293T cell line.

All statistical tests and *P*-values are included in **Supplementary Table 1**.

### Constructs and strains

A derivative of BL21-AI cells was used for all *E. coli* variant library experiments. This derivative, bSLS.114, has the endogenous Retron-Eco1 operon replaced by a chloramphenicol resistance cassette flanked by FRT recombinase sites using the method developed by Datsenko and Wanner^28^. This knock-out cassette was amplified from pKD3, adding homology arms to the Retron-Eco1 locus with PCR primers, and electroporated into BL21-AI cells with the Lambda Red recombination machinery (pKD46). After selecting clones on 10 µg ml^−1^ chloramphenicol plates, we genotyped to confirm the locus-specific knock-out and then excised the chloramphenicol resistance cassette using the FLP recombinase (pMS127).

All yeast variant libraries were cloned into pKDC.100, which contains, under control of a Gal7 promoter, the 5’ end of the *msr/msd* and PaqCI Golden Gate restriction enzyme sites at the 3’ end of the *msd* for insertion of variant parts. This plasmid contains a URA3 selection marker and an episomal origin of replication (CEN/ARS), and was constructed using Gibson assembly, with a Twist-synthesized gBlock containing the PaqCI sites and a PCR-amplified linear pSCL039^15^. Yeast plasmids containing the three editing sites in the HIS3 site were based off pZS.157^14^. These three variants (pSCL194: site 1; pSCL195: site 2; pSCL368: site 3) all contain galactose-inducible Retron-Eco1 RT and *S. pyogenes* Cas9 (Gal1-10 promoter) along with their respective sites. These plasmids were all constructed using Gibson assembly, using pZS.157 to create the backbone and Twist-synthesize gBlocks containing the editing sites. The strains containing these editing sites along with Cas9 and Retron-Eco1 RT were made using LiAc/SS carrier DNA/PEG transformation^29^ of BY4742^30^. The respective plasmids were linearized using KpnI and transformed into BY4742 for homologous recombination into the *HIS3* locus. Clones with selected on SD-HIS media.

All human vectors are derivatives of pSCL.273^5^, itself a derivative of pCAGGS. pCAGGS was modified by replacing the MCS and rb_glob_polyA sequence with an IDT gblock containing inverted BbsI restriction sites and a SpCas9 tracrRNA, using Gibson Assembly. The resulting plasmid, pSCL.273, contains an SV40 ori for plasmid maintenance in HEK293T cells. The strong CAG promoter is followed by the BbsI sites and SpCas9 tracrRNA. BbsI-mediated digestion of pSCL.273 yields a backbone for single or library cloning of plasmids by Gibson Assembly or Golden Gate cloning. Our backbone incorporated an EGFP-P2A and Eco1RT into pSCL.273. Twist-synthesized gBlocks encoding our various ncRNA donors were cloned into this backbone (pKDC.154) via Golden Gate Reaction with PaqCI. Plasmids were subsequently midi-prepped according to manufacturer instruction (Qiagen 12143). Human experiments were carried out in a HEK293T cell line which expresses Cas9 from a Piggybac-integrated, TRE3G-driven, doxycycline-inducible (1 μg/ml) cassette, which we have previously described^15^.

All strains/lines are listed in Supplementary Table 3, and all plasmids in Supplementary Table 2.

### Variant library cloning

*E. coli* variant cloning was done as previously described^15^ using BsaI Type IIS restriction sites and Golden Gate cloning. After high-efficiency cloning and electroporation, variant libraries were miniprepped for electroporation into the experimental strain (bSLS.114, described above). All *E. coli* variant parts were synthesized by Agilent.

All *S. cerevisiae* variant parts were synthesized by Twist. The variant part of the editron ncRNA was flanked by PaqCI Type IIS restriction sites and specific primers to amplify out sublibraries from a larger synthesis run. Each variant part was padded by random nucleotides to 250 bp on the 3’ end, and sublibraries were segregated by original variant part length (gated to each sublibrary having < 10% variance in the length) to avoid library bias with amplifying out sublibraries by PCR. Variant sublibraries were then combined with pKDC.100 in a Golden Gate reaction using PaqCI and the PaqCI activator (2:1 ratio), and T4 DNA ligase (NEB) to generate cloned sublibraries at high efficiency after electroporation into a cloning strain (ECloni Elite 10G, Biosearch Technologies). Sublibraries were then midiprepped and combined based on the number of variant parts in the sublibrary and the DNA concentration to create a final pooled library with equal distribution of variant parts (QIAGEN).

### Variant library expression and sequencing

*E. coli* variant libraries were grown overnight and diluted 1:500 into expression media (arabinose and IPTG for the ncRNA, and erythromycin for the RT). At dilution, we also took a pre-expression sample. We then grew the cells for 5 hours shaking at 37 C. After expression, we took two samples: one for variant plasmid quantification and the other for RT-DNA quantification. The pre-expression and post-expression plasmid samples were mixed 1:1 with water and boiled at 95 C for 5 minutes. The RT-DNA expression sample was prepared as previously described^15^. Briefly, DNA was purified using a modified miniprep protocol, treated with RNase A/T1 (New England Biolabs), and purified with ssDNA/RNA Clean & Concentrator kit from Zymo Research.

After ssDNA isolation, we either amplified the DNA barcode with primers containing Illumina adapters (*msr* sublibraries and plasmid samples) or performed a non-sequence-biased sequencing preparation (*msd* sublibraries). To amplify RT-DNA without prior knowledge of the sequence, we treated the sample with DBR1 (Origene), extended the 3’ end with dCTP with TdT. We used Klenow fragment (3’→5’ exo-) to create the second complementary strand using a primer with six guanines and an Illumina adapter. After creating the second strand, we ligated an Illumina adapter to the 3’ end of the complementary strand using T4 ligase. All products were indexed and sequenced on the Illumina MiSeq. Sequencing primers are listed in Supplementary Table 4.

All yeast variant libraries were transformed into their matched strain using a 40x scaled-up version of the Zymo Frozen-EZ Yeast Transformation II Kit. After a recovery for 1 hour in YPD and an overnight growth shaking at 30 C in 2% raffinose SD -URA -HIS, a time=0 hr sample was taken and then yeast were passaged to 0.2 OD into 50 mL 2% galactose SD -URA -HIS. Cells were then grown for 24 hours shaking at 30 C, a time=24 hr sample was taken. The yeast were then passaged again to 0.2 OD in 50 mL 2% galactose SD -URA -HIS and grown for another 24 hours shaking at 30 C. After a total of 48 hours of editing, the yeast optical densities were measured again and two aliquots of 500 million cells each were collected for the time=48 hours plasmid and genome sample.

Yeast gDNA was extracted as previously described^15^. Briefly, cells were lysed in 120 μL lysis buffer (100 mM EDTA pH 8, 50 mM Tris-HCl pH 8, 2% SDS) and boiled for 15 min at 100 C. After cooling the lysate on ice, proteins were precipitated by adding 60 uL of ice-cold 7.5 M ammonium acetate and incubating at −20 C for 10 min. The samples were centrifuged at 17,000g for 15 min to pellet the protein, and the supernatant containing the gDNA was transferred to a new tube. The gDNA was precipitated in 1:1 ice-cold isopropanol at 4 C for 15 min, and then washed twice with 200 μL ice-cold 70% ethanol. The DNA pellet was dried at 65 C for 5-10 minutes to evaporate all ethanol, and resuspended in 40 μL water. Genomic DNA samples for deep-sequencing were then amplified using primers around the editing site containing Illumina adapters. All products were indexed and sequenced on the Illumina MiSeq. Sequencing primers are listed in Supplementary Table 4.

Yeast plasmid DNA was extracted as previously described^31^. The Zymo Yeast Miniprep Kit was scaled up to 500m cells. Briefly, we resuspended yeast in 1 mL digestion buffer and 30 uL zymolyase, and digested the cell wall for 3 hrs shaking at 900 rpm at 37 C. We then added 1 mL of solution II (lysis buffer) to the tubes, split the sample across multiple microcentrifuge tubes, and added 1:1 solution III (protein precipitation buffer). We then spun down the tubes and sequentially added the supernatant to the Zymo Yeast Miniprep spin column. After reconsolidating the sample, we washed the spin column with 550 μL wash buffer and eluted in 20 μL pre-warmed ultra-pure nuclease-free H2O at 37 C.

To prepare the plasmid samples for sequencing without the creation of hybrid products, we amplified the plasmid barcodes using 50 ng of plasmid DNA and 16 cycles of amplification, performing 8 reactions in parallel per sample using primers containing the Illumina adapters. We then pooled the PCRs for each sample and removed primer-dimers through size-selective bead clean-up. We then use 5 μL of the cleaned-up plasmid DNA amplicons for indexing and sequencing on the Illumina MiSeq. Sequencing primers are listed in Supplementary Table 4.

### Machine learning submethods

We split the retron-Eco1 ncRNA variants and the associated RT-DNA production values into 2930 training sequences, 154 validation sequences, and 342 test sequences. We then trained a convolutional neural network using one-hot-encoded retron ncRNA sequences as inputs and RT-DNA production as the output. The model parameters that were optimized using Ray Tune were number of layers, step size, and number of dilations with a 3:1 train:validation scheme. The final model was made of two computational blocks and a residual dilated convolution block followed by a two-layer perceptron. All model code will be available on GitHub prior to peer-reviewed publication.

### Human editing expression and analysis

All HEK cells were cultured in DMEM + GlutaMax supplement (ThermoFisher 10566016) + 10% HI-FBS. 6-well cultures were transiently transfected with 7.32 ug of plasmid per well using Lipofectamine 3000 (ThermoFisher). 24 hrs after transfection, doxycycline was refreshed and cultures were passaged into T-25 flasks to be grown for an additional 48 hrs. Three days after transfection, cells were collected for FACS sorting. DAPI dye was added to stain for live/dead and cells were gaited on DAPI and GFP with untransfected cells used as a negative control for background (BD FACSAria Fusion).

### Human sample preparation

To prepare samples for sequencing, sorted cells were collected and gDNA was extracted using a QIAamp DNA Mini Kit according to the manufacturer’s instructions. DNA was eluted in 50 μl of ultra-pure, nuclease-free water.

2 μl of the gDNA was used as template in 25-μl PCR reactions with primer pairs to amplify the locus of interest which also contained adapters for Illumina sequencing preparation. Lastly, the amplicons were indexed, and sequenced on an Illumina MiSeq/NextSeq instrument.

### RT-DNA production quantification

RT-DNA production was quantified as previously described^15^. Briefly, custom Python software was used to extract the variant counts from the plasmid and RT-DNA samples. We then normalized raw counts to relative abundance (raw count over the total number of raw counts) and RT-DNA relative abundance to plasmid relative abundance, using the average of the pre- and post-induction plasmid abundances to integrate the plasmid abundance over the 5-hr expression window. Finally, these relative abundances were normalized to the retron-Eco1 wild-type abundance, set at 100%.

### Editing rate quantification

Custom software was built to quantify library-scale and individual validation editing rates in yeast and human cells. For yeast variant libraries, raw barcode counts were pulled from the 48-hr genome (editing site) samples, and the 0-, 24-, and 48-hr plasmid samples. The read counts from the plasmids were summed across the three time samples to integrate the plasmid abundances over the editing window, and then each barcode read count was normalized against all barcode read counts in that sample. The relative abundance of an editor’s barcode in the genome was then divided by the relative abundance of an editor’s barcode in the integrated plasmid pool.

For human validation of individual variants, custom software was used to assess the number of reads with the precise edit divided by the number of reads with the wild-type sequence. All software used in the analysis of this paper is available on GitHub.

## RESULTS

### RT-DNA Production in *E. coli* from Retron -Eco1 ncRNA Variant Libraries

The retron-Eco1 ncRNA is a highly-structured RNA molecule with characteristic stem-loops and double-stranded regions that is partially reverse transcribed to generate abundant RT-DNA in cells (**Fig 1a**). As previous work found RT-DNA production was a limiting factor in using the retron as a template for precise editing in prokaryotes and eukaryotes and as a DNA barcode for molecular recording^5,18,18^, we set out to systematically understand how variations in ncRNA sequence and structure impact RT-DNA production in *E. coli*. We constructed a 3,443 member library of ncRNA variants, changing both the *msr* (non-reverse transcribed region) and *msd* (reverse transcribed region). This library contained all single-nucleotide substitutions, scanning deletions and insertions of varying sizes, and variations on length and complementary of stem-loops and all permutation of the three-nucleotide RT recognition motif in the P3 loop. For variants with changes in the *msr*, we included a linked barcode in the P4 loop of the RT-DNA to allow amplification of the barcode via PCR. The library was constructed using Golden Gate cloning, transformed into a B-strain *E. coli* bSLS.114 (BL21-AI Δretron-Eco1), and expressed along with the retron-Eco1 RT for 5 hours, after which we collected RT-DNA for quantification. All variant sequences are included in **Supplementary Table 9-18**.

**Figure 1.**
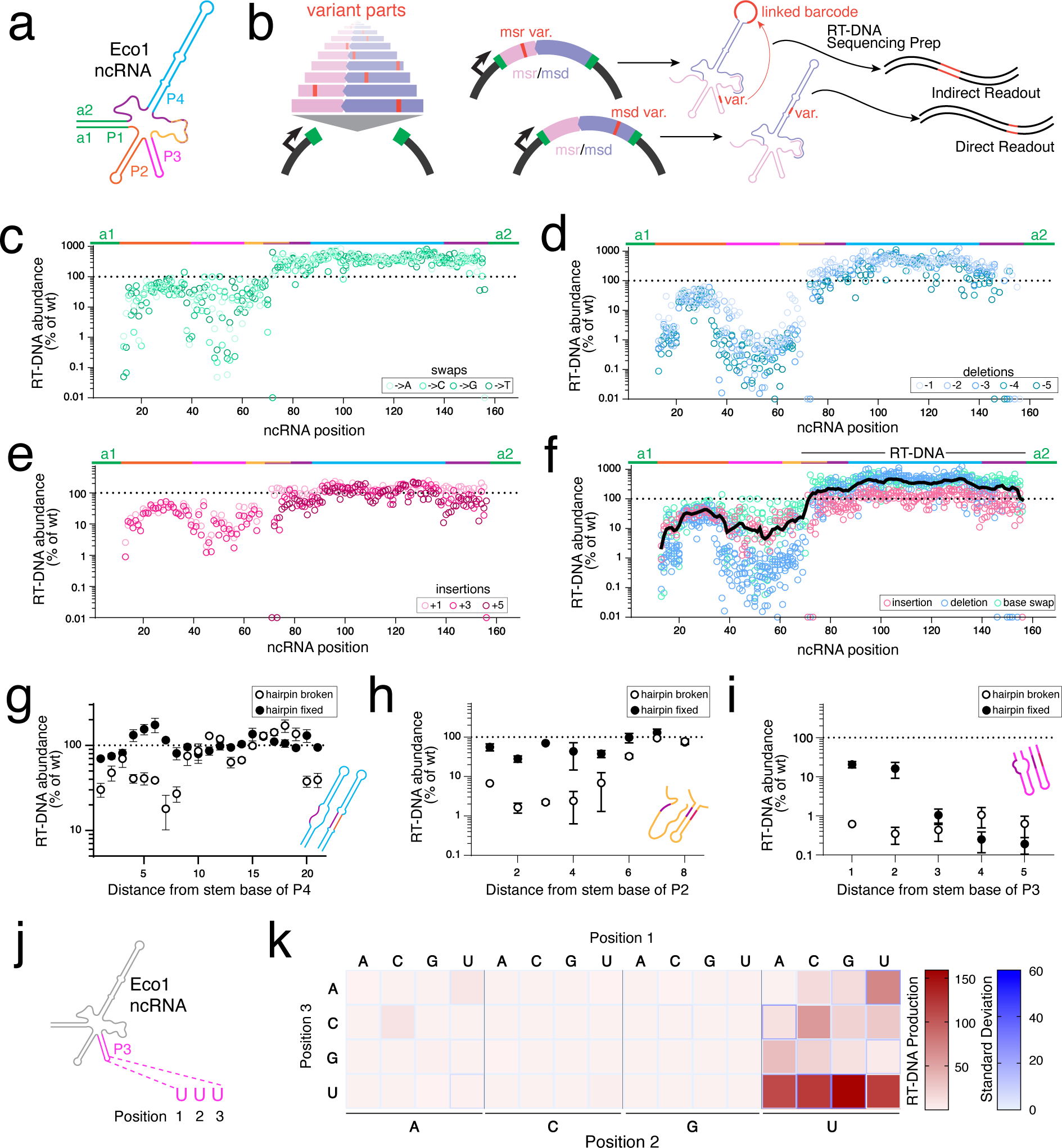
RT-DNA production of retron -Eco1 variant libraries in *E. coli*. **a.** Wild-type -Eco1 ncRNA structure. **b.** Variant library schematic: variants were introduced on the *msr* (non-reverse transcribed part of the ncRNA) or the *msd* (reverse-transcribed part of the ncRNA). After production of the RT-DNA libraries in *E. coli,* single-stranded DNA was sequenced and variants quantified. *msd* variants were identified on the RT-DNA, while *msr* variants were identified through a barcode in the P4 loop. **c.** RT-DNA production of all single-nucleotide substitutions relative to wild-type RT-DNA. Each open circle represents the mean of three biological replicates. **d.** RT-DNA production of 1, 2, 3, 4, and 5 nucleotide deletions starting at a specified ncRNA position relative to wild-type RT-DNA. Each open circle represents the mean of three biological replicates. **e.** RT-DNA production of 1, 3 and 5 nucleotide insertions starting at a specified ncRNA position relative to wild-type RT-DNA. Each open circle represents the mean of three biological replicates. **f.** Summary of RT-DNA production relative to wild-type RT-DNA production of all single-nucleotide variants: insertions (pink), deletions (blue), and substitutions (green). RT-DNA production relative to wild-type RT-DNA is shown across the nucleotide positions in the ncRNA from 5’ to 3’. The black line on top is the mean of RT-DNA production of all the changes at that nucleotide position. Each open circle represents the mean of three biological replicates. **g.** RT-DNA abundance of removing complementarity (black) and restoring complementarity (white) of stem P4 with different nucleotides along the distance from stem base relative to wild-type RT-DNA abundance. Each circle represents the mean of three biological replicates with error bars representing the standard error. The effect of breaking the stem is significant (one-way ANOVA using only broken stem and wild-type data, *P*<0.0001) at positions 1, 4, 5, 6, 7, 8, 18, 20, and 21 compared to the wild-type stem (position 1, *P*=0.005; position 4, *P*=0.0254; position 5, *P*=0.0261 position 6, *P*=0.0194; position 7, *P*=0.0007; position 8, *P*=0.003; position 18, P=0.0045; position 20, *P*=0.0164; position 21, *P=*0.0208) (Dunnett’s, corrected). Restoring the stem structure significantly increases RT-DNA production only at positions 7 and 21 (position 7, *P=*0.0023; position 21, *P*=0.0285) (Bonferroni corrected for multiple comparisons). **h.** RT-DNA abundance of removing complementarity (black) and restoring complementarity (white) of stem P2 with different nucleotides along the distance from stem base relative to wild-type RT-DNA abundance. Each circle represents the mean of three biological replicates with error bars representing the standard error. The effect of breaking the stem is significant (one-way ANOVA using only broken stem and wild-type data, *P*<0.0001) at all positions compared to the wild-type stem except position 7 compared to the wild-type stem (position 1, *P*<0.0001; position 2, *P*<0.0001; position 3, *P*<0.0001; position 4, *P*<0.0001; position 5, *P*<0.0001; position 6, *P*<0.0001; position 7, P=0.7977; position 8, *P*=0.0029) (Dunnett’s, corrected). Restoring the stem structure significantly increases RT-DNA production at positions 1, 2, 3, and 5 (position 1, *P=*0.01; position 2, *P=*0.001; position 3, *P*<0.0001; position 5, *P=*0.03) (Bonferroni corrected for multiple comparisons). **i.** RT-DNA abundance of removing complementarity (black) and restoring complementarity (white) of stem P3 with different nucleotides along the distance from stem base relative to wild-type RT-DNA abundance. Each circle represents the mean of three biological replicates with error bars representing the standard error. The effect of breaking the stem is significant (one-way ANOVA using only broken stem data, P<0.0001) at all positions compared to the wild-type stem (position 1, *P*<0.0001; position 2, *P*<0.0001; position 3, *P*<0.0001; position 4, *P*<0.0001; position 5, *P*<0.0001) (Dunnett’s, corrected). Restoring the stem structure only significantly increases RT-DNA production in position 1 (*P=*0.0041) (Bonferroni corrected for multiple comparisons). **j.** Eco1 reverse transcriptase recognition motif UUU in the terminal loop of stem P3. **k.** RT-DNA production of every permutation of retron-Eco1 reverse transcriptase recognition motif relative to wild-type RT-DNA abundance. Position 1 is shown at the top of the heat map, Position 3 on the left, and Position 2 on the bottom. RT-DNA production is scaled with on the red-white colorbar, while the standard deviation is represented by the blue around the squares of the heatmap. Each square represents the mean of three biological replicates. There is a significant effect of the RT recognition motif (one-way ANOVA, *P*<0.0001), with every permutation significantly different than the wild-type UUU (*P*<0.0001) except UUA and AUU (*P*=0.8991 and *P*=0.0551, respectively) (Dunnett’s, corrected).

To quantify the RT-DNA abundance of *msd* variants, we used a sequencing pipeline described previously that allows us to amplify RT-DNA without requiring prior knowledge of the RT-DNA sequence^5,15,23^. Briefly, we (1) purified short single-stranded DNA (ssDNA) using a QIAGEN Midiprep Plasmid Plus Kit followed by a Zymo ssDNA Clean & Concentrator Kit, (2) treated the resulting ssDNA with Dbr1 to remove the 2’-5’ linkage between the RT-DNA and ncRNA, (3) extended the debranched ssDNA with a single polynucleotide using template-independent polymerase (TdT), (4) generated a complementary strand using a primer consisting of the complementary single polynucleotide and an Illumina adaptor, (5) ligated an adaptor to the other end of the now double-stranded RT-DNA, and lastly (6) Illumina sequenced the now double-stranded RT-DNA with Illumina adaptors on both ends. RT-DNA barcodes linked to changes in the *msr* were quantified by amplifying the barcode for sequencing after purifying ssDNA. All variants were normalized against the production of the wild-type retron-derived RT-DNA and the abundance of the variant plasmid (**Fig 1b**). To quantify the relative abundance of each variant plasmid in the expression cells, we amplified the variable region of the ncRNA using plasmid-specific primers and sequenced the amplicons using Illumina sequencing.

Figure 1c shows single-nucleotide substitutions scanning across the retron-Eco1 ncRNA, where we found substantial sequence flexibility on the single-nucleotide level except at two important positions: around the priming guanosine immediately after the a1 region, important for making the 2’-5’ linkage of ncRNA-to-RT-DNA; and around the UUU putative recognition loop for the retron-Eco1 reverse transcriptase (**Fig 1c, Supplementary Figs 1b-e**).

We also analyzed deletions scanning across the retron-Eco1 ncRNA that varied in length from 1 to 5 nucleotides. The retron-Eco1 ncRNA is less tolerant to deletions than substitutions, particularly in the *msr* P2 and P3 stem loops, suggesting a greater influence of structure over sequence. In addition, deletions in the *msd* region directly flanking the a2 region were not tolerated. Larger deletions are less tolerated than smaller deletions in the critical region of the P3 stem-loop (**Fig 1d, Supplementary Figs 1f-j**).

We also assessed 1-, 3-, and 5-nucleotide insertions scanning across the retron-Eco1 ncRNA (**Fig 1e**). While small insertions were slightly more tolerated than small deletions (**Supplementary Fig 1g, Supplementary Figs 1k-l**), larger insertions in the *msr* region resulted in undetectable levels of RT-DNA (**Supplementary Fig 1m**). Similarly to deletions, insertions directly adjacent to the a2 region in *msd* also greatly reduced RT-DNA production.

A summary of the effect of all nucleotide substitutions, deletions, and insertions is shown in **Fig 1f**. Generally, the retron-Eco1 ncRNA tolerates modifications in the P2 and P4 stem-loops, but is relatively intolerant to modifications around the priming guanosine and the stem-loop P3. The tolerance to mutations in the P3 stem is important for the use of retron-Eco1 in biotechnology, as this is the position where editing donors and DNA barcodes have been encoded.

Next, we sought to assess the effect of structural variations. To do this, we quantified the effect of breaking complementarity in stem-loops P2, P3, and P4 by replacing one side of the stem with a non-complementary new sequence to create a nucleotide bubble of length 4 in stem-loops P2 and P3, and length 5 in stem-loop P4. To control for an effect of a sequence versus structural change, we also restored complementarity by changing the same position on the other side of the stem with the complement of the replaced nucleotides. Breaking P4 complementarity only affected RT-DNA production at the base and the tip of the stem, and fixing complementarity with different sequences restored wild-type levels of RT-DNA production (**Fig 1g**). Breaking stem-loop P2 complementarity closer to the base reduced RT-DNA production and restoring complementarity restored RT-DNA production (**Fig 1h**). Breaking stem-loop P3 complementarity reduces RT-DNA production, but restoring complementarity with an alternate sequence does not restore RT-DNA production (**Fig 1h**). Overall, there are clear structural requirements, most notably in P2/3: in P2, structure is important; and in P3, both sequence and structure are important for RT-DNA production.

We next sought to quantify how strictly required the UUU recognition motif in the loop of P3 is for reverse transcriptase recognition (**Fig 1j**). Testing every permutation of the UUU motif reveals low sequence flexibility in position 3 (vertical axis) and position 2 (bottom axis), requiring both of these to be uracils. However, there is significant flexibility in position 1, with every possible base approaching wild-type RT-DNA production levels (lower right square, UUU), with GUU having higher RT-DNA production than wild-type (**Fig 1k**).

### Machine Learning on Libraries Reveals Novel Variables to Increase RT-DNA Production

Though we tested ∼3,400 variants of the retron-Eco1 ncRNA including all single-nucleotide substitutions, a variant library of all possible nucleotide combinations would number on the order of 10^90^ variants, without including insertions and deletions. Therefore, to explore more of the possible sequence space, we used the ncRNA variant library data to create a machine learning algorithm capable of predicting novel retron ncRNA sequences with enhanced RT-DNA production. The experimental values across ∼3400 measurements were inverse normal transformed and split into a train, validation, and test sets. A convolutional neural network, named retDNN, was then used to learn the relationship between sequence and RT-DNA levels. The retDNN model comprises of two computational blocks and a residual dilated convolution block followed by a two-layer perceptron. The model was trained on 3084 measurements and tested on the held-out set, achieving an *R*=0.671 performance (*R*=0.775 on the training set) (**Fig 2a-b**). We then queried the retDNN model with *in silico* variants, including a P4 stem-loop of varying GC content. Interestingly, the model predicted that lowering GC content in the P4 stem-loop would increase RT-DNA production over wild-type, something untested in the original variant library. To validate this prediction, we synthesized and cloned the 500 queried variants of differing GC contents (25 variants per 10% GC content range) and experimentally validated RT-DNA production relative to wild-type through the same sequencing pipeline as above. As the algorithm predicted, lower GC percentages of the P4 stem-loop produced more RT-DNA (**Fig 2c**).

**Figure 2.**
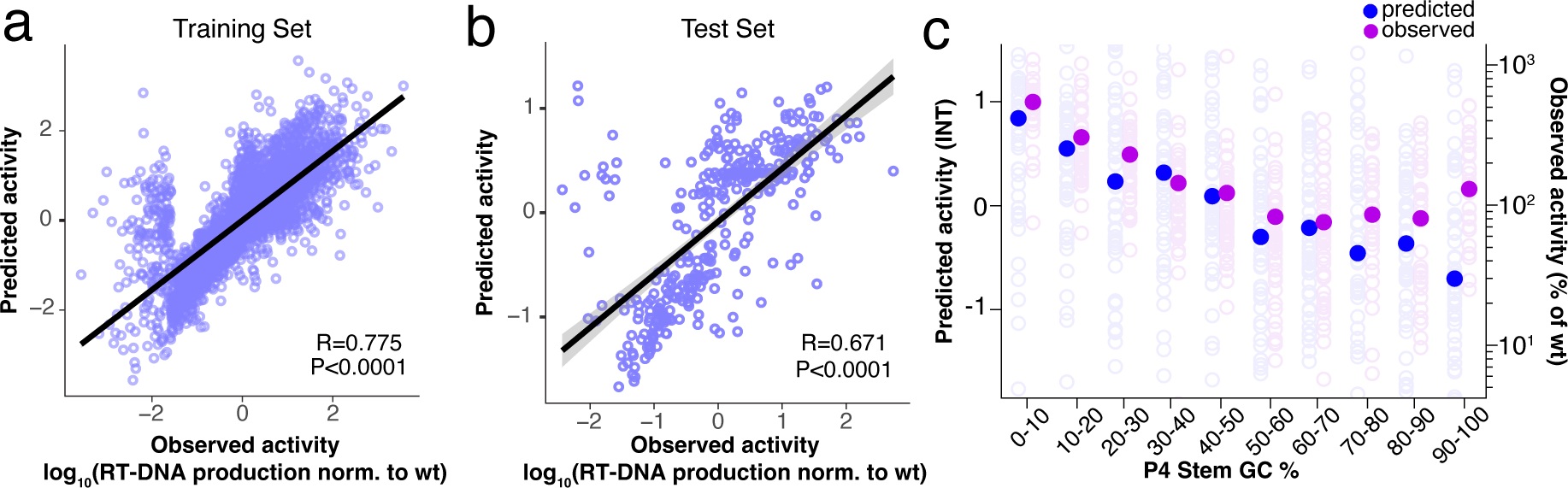
Machine learning on variant libraries guides novel predictors of RT-DNA production. **a.** Machine learning algorithm performance on training set of ncRNA variants from *E. coli*. Input is ncRNA sequence and output is inverse-normalized variant RT-DNA production. Each open circle represents an individual ncRNA sequence. Linear regression *R* and *P*-values of ML predicted activity vs. observed activity annotated on the plot. **b.** Machine learning algorithm performance on held-out test data. Each open circle represents and individual ncRNA sequence. Linear regression *R* and *P*-values of ML predicted activity vs. observed activity annotated on the plot. **c.** Predicted (blue) and experimentally-determined (purple) RT-DNA production of varying GC percentages in stem P4. Open circles represent means of two biological replicates of individual ncRNA variants and closed circles represent the mean of all ncRNA variants tested for that GC percentage. Linear regression slope of the predicted (blue) points has a slope of −0.0156 and a *P*-value of <0.0001. Linear regression slope of the observed (purple) points has a slope of −3.7995 and a *P-*value=0.0069.

### Editing Performance in *S. cerevisiae* of Retron-Eco1 ncRNA Variant Libraries

Efficient RT-DNA production is critical for retron biotechnology, including the use of RT-DNA as the donor for precise genome editing. In this context, a retron-Eco1 ncRNA is modified to encode a precise repair donor in the stem-loop of P4 and a guide RNA for Cas9 double-strand DNA cleavage at the 3’ end of the ncRNA. This combination of CRISPR-Cas9 and retron immune systems has been called CRISPEY in yeast^14^ or as an editron to encompass its use in eukaryotic cells. After determining the effect of ncRNA variations on RT-DNA production in *E. coli*, we sought to extend this understanding to editing and additionally investigate how donor, guide RNA, and ncRNA chassis variants all together affect precise editing rates in eukaryotes.

We designed a library to assess the contributions of structural, cut site, and donor variables to precise genome editing, with each variant inserting a unique 10-bp barcode into the yeast genome at a designated site, along with changing the NGG *S. pyogenes* Cas9 protospacer adjacent motif (PAM) to NAT to prevent re-targeting of the edited site. We synthesized variant libraries for the same variables across three unique sites: two artificial, constructed sites with designed, symmetric PAMs around the edit site, and one site from the human genome (an intron in the *NPAS2* gene) with the same PAM locations as the constructed sites. These three sites were independently integrated into the *HIS* locus of *S. cerevisiae* to interrogate the local sequence effects on the editing efficiency, while ensuring the editing site remains active and open by also providing a copy of the *HIS* gene in *HIS* auxotrophic yeast, and maintaining strains in -HIS media.

In these variant libraries, we assessed: 5 donor lengths (54, 64, 78, 94, and 112 nucleotides), 5 homology arm symmetries about the edit site per donor length, RT-DNA donors that are complementary to the target or non-target strand, and 5 different cut sites (−16, −8, 0, +8, +16 relative to barcode insertion point), leading to 175 donor/gRNA combinations per site (**Fig 3a**). We then combinatorially combined these donor/gRNA variants with 25 different ncRNA chassis: wild-type -Eco1 ncRNA, CRISPEY ncRNA^14^, the 13 best-performing structural variants from the *E. coli* variant libraries, and 10 *de novo* predicted ncRNAs from the machine learning algorithm. In all, we tested 3,125 variants per site. All variant sequences are included in **Supplementary Table 6-8**.

**Figure 3.**
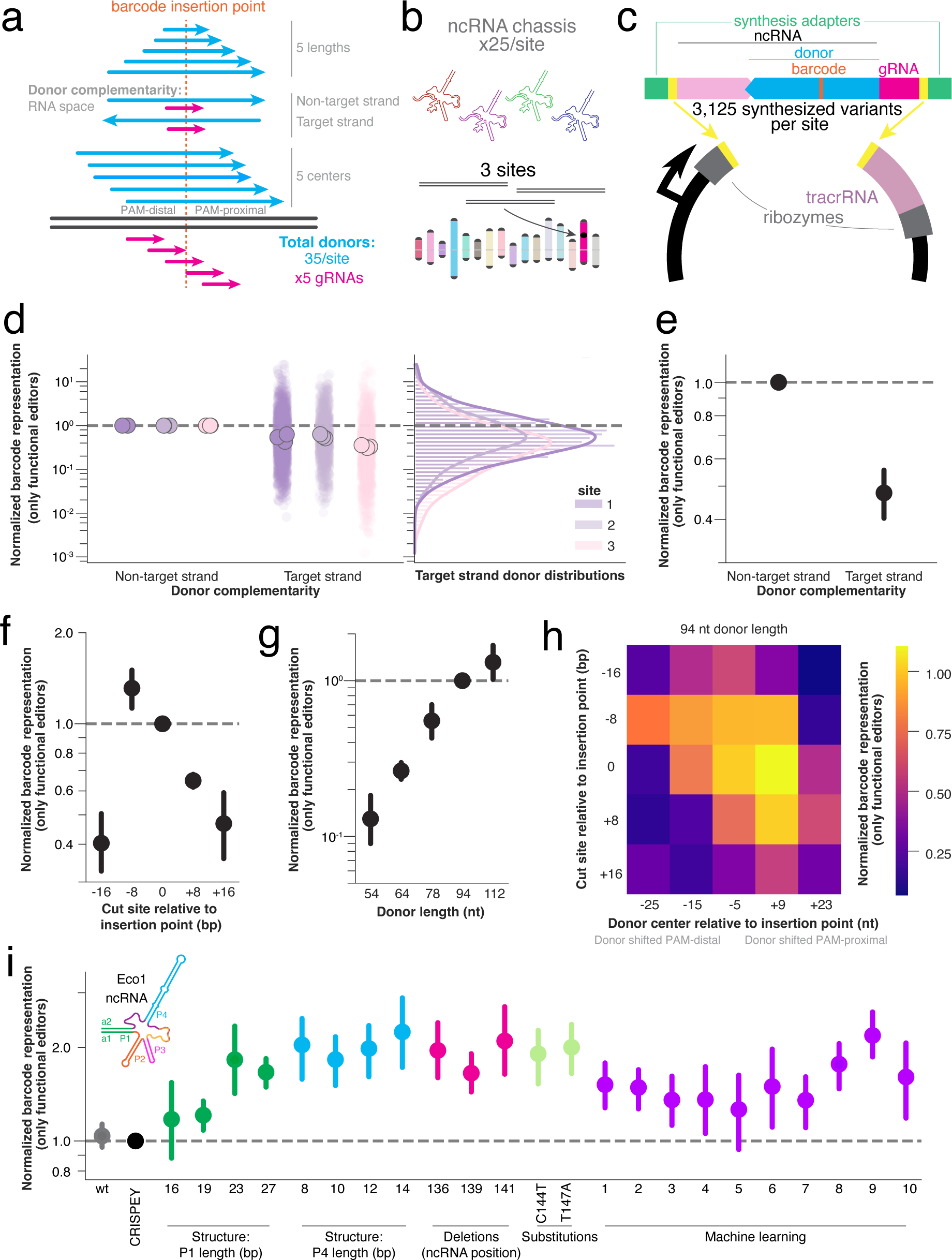
Precise editing of retron Eco1 editing variant libraries in *S. cerevisiae*. **a.** HDR donor variant schematics and gRNA variants, with 5 donor lengths, 2 donor directions relative to the gRNA, and 5 donor centers relative to edit and cut position for a total of 50 donors per editing site. There are 5 evenly-spaced gRNAs per site relative to the edit position, for 250 donor/gRNA pairs per site. **b.** There are 25 ncRNA chassis per donor/gRNA combination. **c.** Three sites integrated into the HIS locus of the yeast genome were tested: two synthesized and one from the human genome (NPAS2 locus). **d.** Schematic for 3,125 variant plasmids per site in the library. Each variant has a unique 10 bp barcode that can be read out from the plasmid or from the edit site in the genome. **e.** All complementary gRNA/donor variants’ barcode representation normalized against its parallel gRNA/donor variant, with all other variables held constant (chassis, donor length, center, and gRNA). The variants for each site are plotted in different colors, and each biological replicate of a site is summarized by the median (left) of the distribution of variants (right). **f.** Data in Fig 3e summarized as the mean of all sites and all biological replicates (closed circle) (±standard deviation), with target-strand-homologous donors editing at significantly lower frequencies (1-sample t-test; *P*<0.0001). g. Barcode representation of cut sites normalized to the cut site at the barcode insertion site (±standard deviation), with cut sites at −16, +8, and +16 editing at significantly lower frequencies (1-sample t-test, Bonferroni correction for multiple comparisons; *P*<0.0001, *P*<0.0001, *P*<0.0001 respectively, all other comparisons non-significant). **h.** Barcode representation of donor lengths normalized to 94 nucleotide donor length (±standard deviation), with donor lengths <94 nucleotides editing at significantly lower frequencies (1-sample t-test, Bonferroni correction for multiple comparisons; *P*<0.0001, *P*<0.0001, *P*<0.01 respectively, all other comparisons non-significant). **i.** Heat map of normalized barcode representation of cut site vs. donor center (94 nucleotide donor length), normalized to the cut site at the barcode insertion site, and donor center of 5 bp upstream the barcode insertion site. Cut site and donor center interact significantly (two-way ANOVA; *P*-value of interaction < 0.0001) **j.** Barcode representation of all chassis ncRNA normalized to the CRISPEY ncRNA (±standard deviation) Chassis with a1/a2 27-bp length, 10-bp and 12-bp P4 length, deletion at position 139, substitutions at C144T and T147A, and ML chassis 8 and 9 all edit at significantly higher frequencies (1-sample t-test, Bonferroni correction for multiple comparisons; *P*=0.004, *P*=0.028, *P*=0.036, *P*=0.019, *P*=0.049, *P*=0.019, *P*=0.024, and *P*=0.009 respectively).

Three independent yeast lines were created, each with one of the three sites in the *HIS* locus of the yeast genome along with Cas9 and retron-Eco1 RT under the control of a *GAL1/10* galactose-inducible divergent promoter (**Fig 3b**). These synthesized ncRNA variants for each site were encoded on a vector containing other necessary ncRNA components (ribozymes, tracrRNA) under the *GAL7* galactose-inducible promoter (**Fig 3c**). After transformation of the editing libraries into yeast, editing was performed for 48 hours in galactose media.

To analyze the data, we sequenced the barcode distribution in the plasmid pool and the barcodes inserted into the correct site in the yeast genome after 48 hours of editing. First, we calculated the proportion of each barcode’s reads in the pool of reads (for barcodes edited into the genome: the reads at 48 hours of editing; for barcodes in the plasmid pool: the reads as summed over samples taken at 0, 24, and 48 hours of editing). This is to integrate the plasmid barcode pool over the entire editing period. Plasmid barcode read count was stable over the 48 hours of editing (**Supplementary Fig 2**). Then, we normalized the individual barcode proportions as seen in the genome to the same barcode’s proportion as seen in the plasmid pool (called barcode representation henceforth), and removed barcodes not seen at counts >10 in the plasmid pool or not seen at all in the genome pool (percent of working editors per library variable is shown in the Supplementary Figures). We then normalized along the axis of interest. For example, when assessing the effect of donor RT-DNA complementary to either the target or non-target strand (target strand: strand complementary to the gRNA/complementary to the PAM-containing strand; non-target strand: strand not complementary to gRNA/PAM-containing strand), we held all other variables constant (donor length, cut site, donor center, chassis) and normalized the target strand barcode representation to the non-target strand barcode representation of each specific group. This normalized barcode representation for every barcode for each biological replicate for each site is represented as a transparent circle in **Fig 3d**. We then took the median of each biological replicate of each site, based on the distribution on the right of **Fig 3d**, and averaged those across all sites to obtain the summary figure for that axis of interest. After performing this normalization, we found that, on average, target strand donors are worse editors than non-target strand donors because the barcode was inserted less often when holding all other variables constant, performing at about 50% efficiency as compared to the matched non-target strand donors (**Fig 3e**). Both target strand and non-target strand donors have about 50% functional editor variants, as other parameters also influence if an editor is functional (**Supplementary Fig 3a**).

We analyzed the effect of cut site positioning relative to insertion point by using Cas9 spacer sequences 8 nucleotides apart and analyzing as above, normalizing within-group to a cut position of 0, the site at which Cas9 cuts directly where the 10-bp barcode is then inserted. We noticed that the cut site of −8 for site 2 had an unusually low number of working donors (<20%), indicating that this guide RNA has low cutting efficiency, and so excluded that cut site for site 2 from the analysis (**Supplementary Fig 3b**). We found that an edit on the PAM-proximal side of the cut site performed slightly better (∼130% efficiency at the cut site of −8 compared to a cut position of 0) and performed much better than an edit on the PAM-distal side of the cut site (∼65% efficiency at the cut site of +8), with consistency across sites, while cut sites far from the insertion point resulted in lower frequency of precise editing (∼40-45% efficiency) (**Fig 3f**). However, it should be noted that only donors complementary to the non-target strand were included in this part of the editron library.

We examined the effect of donor length, normalizing within-group to a donor length of 94 nucleotides. In general, longer donors were more efficient editors than shorter donors, with a 54 nucleotide donor editing at ∼10% of the rate of to the 94 nucleotide donors, while 112 nucleotide had ∼130% efficiency compared to the 94 nucleotide donor (**Fig 3g**). The percentage of working donors per donor length also increased with donor length (**Supplementary Fig 3c**).

We assessed the effect of donor center and cut site together by adjusting the length of the homology arms, normalizing within-group to the centered cut site (0) and centered donor (−5). The data for 94 nucleotide donor length is shown, as each different donor length has different donor center points. All other donor length results are shown in **Supplementary Fig 4**. As the higher normalized barcode representation goes from top left to bottom right, it was generally better to center the donor around the cut site than the insertion point, except for cases of cut sites very far from insertion point (top left and bottom right). In addition, when cut and insertion points were overlapping, a slightly PAM-proximal shifted donor performed slightly better than centered, at 110% efficiency compared to centered (**Fig 3h**). The percentage of working donors for donor center and cut site is included in **Supplementary Fig 5**.

Finally, we analyzed the effect of ncRNA chassis, normalizing within-group to the original CRISPEY chassis. In general, no structural variants performed worse than the CRISPEY chassis, and several variants performed significantly better (27 bp a1/a2 extension, 10 and 12 bp P4 stem length, deletion at position 139, C144T and T147A, and ML chassis 8 and 9) (**Fig 3i**). Excitingly, we found that the machine learning-predicted chassis supported equally high rates of editing despite deviating from the natural sequence by 55-80% in the 20 nucleotide ML variable region, or up to 12% over the full retron-Eco1 ncRNA including the 27-bp extended a1/a2 (logo map of ribonucleotide usage across the machine learning variable region in **Supplementary Figure 7**). Specific machine learning chassis structures and sequences can be found in **Supplementary Figure 6**. We found no evidence of a difference in the percentage of working donors across ncRNA chassis (**Supplementary Figure 8**).

### Library-Informed Optimization of Human Editing

We next sought to understand if design rules learned in *E. coli* and *S. cerevisiae* extend to editing in human cells. Editrons contain the same constituent parts in human cells as in yeast, except for the editing ncRNA is driven by an H1 promoter for nuclear retention rather than being flanked by ribozymes. Our plasmids included an EGFP and retron-Eco1 RT separated by a P2A driven by a CAG promoter and a ncRNA containing an editing donor fused to a sgRNA driven by a Pol III H1 promoter. The addition of the EGFP enables selection of cells successfully transfected with the plasmid. The editing donors consist of homology arms, a PAM recode (NGG>NAT), and a single nucleotide change. We chose to target an intron in the endogenous *NPAS2* site for human validation, using the exact ncRNA constructs used in the yeast libraries. All donors tested are included in **Supplementary Table 5**.

The editron plasmids were transfected into HEK293T cells containing an integrated doxycycline-inducible Cas9, whose expression was induced 24 hours before transfection. Cells were collected three days after transfection and sorted via FACS to only include live and transfected cells, eliminating any variability to due transfection efficiency (**Fig 4a**). We used gRNA 5 for human validation, after an initial screen for gRNA efficacy showed it to have the highest rates of insertions/deletions (indels) of the 3 tested gRNAs, indicating highest cutting efficiency (**Fig 4b**). Consistent with our earlier findings in yeast, we demonstrated that a longer donor and a donor homologous to the non-target strand improve editing efficiency (**Fig 4c-d**). A 112 nucleotide donor increased precise editing from ∼5% to ∼12%, while a non-target strand homologous donor increased editing from ∼4% to ∼12%.

**Figure 4.**
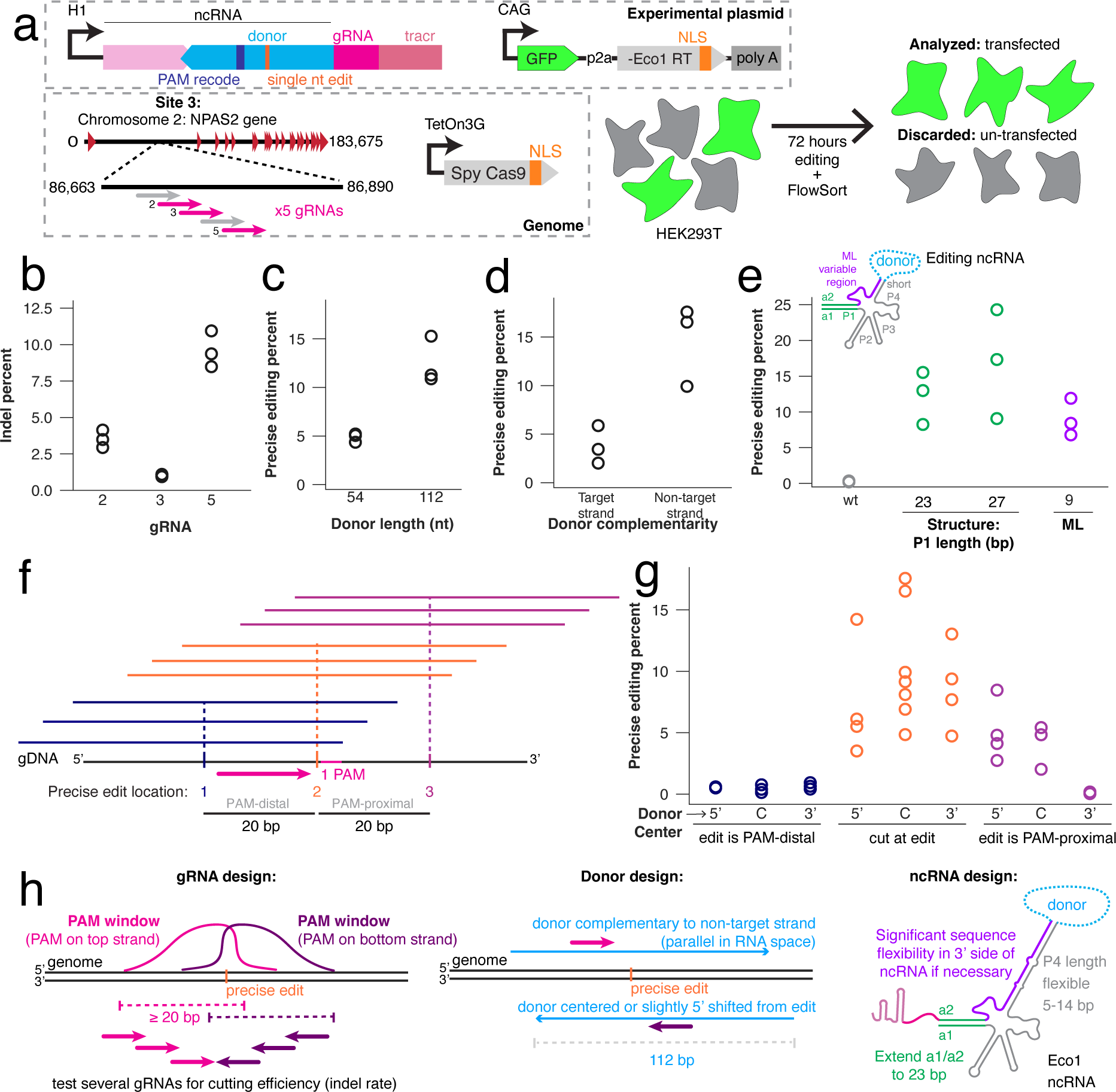
Validating yeast editing libraries with individual human variants. **a.** Human editing schematic. HEK293T cells were transfected with a plasmid containing the editing ncRNA variant with a single nucleotide transversion as a precise edit, along with recoding the PAM NGG to NAT. The plasmid also contained a constitutively-driven GFP-P2A-Eco1 RT. The editron targeted an intronic region of the NPAS2 gene on Chromosome 2 (“site 3” in the yeast data in Figure 3). The HEK293T line also had semi-randomly-integrated *S. pyogenes* Cas9 by PiggyBac transposase under a dox-inducible promoter and a C-terminal NLS. 72 hours after transfection, the HEK293T cells were sorted as GFP+/DAPI-(alive transfected cells) and their genomes were sequenced for precise edits. **b.** Indel percent of the three tested gRNAs. Individual biological replicates are open circles. All gRNA indel rates are statistically different from one another (one-way ANOVA, *P*<0.0001; Bonferroni post-hoc test showed *P*<0.05 for all comparisons). **c.** Precise editing percentages of 52 nucleotide and 112 nucleotide long donors. Individual biological replicates are open circles. The 112 nucleotide donor is a significantly more efficient editor (paired t-test, *P*=0.025). **d.** Precise editing percentages of target and non-target strand homologous donors. Individual biological replicates are open circles. Non-target strand homologous donors are significantly more efficient editors (paired t-test, *P*=0.043). **e.** Precise editing percentages of four ncRNA chassis: wild-type Eco1 ncRNA, extended P1 (a1/a2) (23 and 27 bp), and machine learning chassis 9. Individual biological replicates are open circles. There is a significant effect of ncRNA chassis (one-way ANOVA, *P=*0.01), with a1/a2 extensions of 23 (*P=*0.0267) and 27 bp (*P=*0.0046) performing significantly better than wild-type, and ML chassis 9 not performing worse than wild-type (*P=*0.0993) (Dunnett’s, corrected). **f.** Schematic of donor center relative to precise edit site and cut site. Three precise edits were spaced 20 bp apart, with the cut site centered on the middle edit. Three different donor positions were used per edit: 5’-sided, centered, and 3’-sided. **g.** Precise editing percentages of the 9 different donor center/edit combinations. Three datapoints in the central cut/centered donor are repeated from (d), as these replicates served as the controls for both the donor center/cut site experiment and the target strand experiment. There is a significant effect of edit site and donor symmetry (one-way ANOVA, *P=*0.0002), with all edits on the PAM-distal side of the cut (*P=*0.0014 for 5’ donor center, *P=*0.0012 for centered donor, *P=*0.0016 for 3’ centered donor) and the 3’ donor center on the PAM-proximal side (*P=*0.0009) performing significantly worse than a central cut and edit (Dunnett’s, corrected). **h.** Schematic illustrating final recommendations for editron design.

We chose to validate three chassis modifications in human cells. Longer a1/2 length increased editing compared to wild-type a1/a2 length. Excitingly, ML modifications enabled successful editing despite only 30% sequence similarity to wild-type, demonstrating the flexibility of the region (**Fig 4e**). Next, we sought to determine the ideal positioning of both the edit and the donor relative to a set cut site. We tested 3 edits: a middle edit at the cut site, an edit 20 bp upstream of the cut site, and an edit 20 bp downstream of the cut site. For each of these edits, we tested a donor which was non-symmetric about the edit with more homology on the 5’ side of the non-target strand, centered on the edit, or non-symmetric about the edit with more homology to the 3’ side of the edit site on the non-target strand (**Fig 4f**). All donors used were complementary to the non-target strand. We found that placing an edit at the cut site and on the PAM-proximal side both allowed successful editing, with a slight trend favoring the central cut. Additionally, the trend shows that a donor centered on the cut or with more homology on the PAM-proximal side donor both enable editing. None of the conditions with the edit on the PAM-distal side were edited successfully (**Fig 4g**).

Based on all our variant testing, we provide a set of generalizable design principles for creating future editrons for new targets. Testing several gRNAs to achieve optimal cutting efficiency is an important first step based on our findings showing the variability in indel rates among guides. Donors should be parallel to the guide and complementary to the non-target strand as RT-DNA, with a 112 nucleotide donor having the highest precise editing rate. Additionally, the cut should be centered or non-symmetrically shifted towards the PAM-proximal side of the non-target strand. When modifying the ncRNA, the a1/2 should be extended at least to 23bp. We also demonstrate flexibility in the 3’ region and the P4 length of the ncRNA, allowing for modifications as needed (**Fig. 4h**).

## DISCUSSION

In this work, we comprehensively evaluated the effect of ncRNA variations on RT-DNA production in bacteria from which we trained and validated a ML model. We then evaluated the effect of variations in donor and gRNA, along with ncRNA structure, on editing efficiency in yeast; and validated the major findings in human cells. From these variant libraries, we found that the *msd* region of the ncRNA is generally tolerant to alterations, specifically the stem-loop P4, in which programmable sequences for biotechnology can be inserted, like a donor sequence for precise editing or a transcription factor motif for attenuating transcription factor activity. We also found regions of the *msr* that are required for efficient reverse transcription, such as the stem-loop P3 where the retron-Eco1 RT initiates reverse transcription. In terms of editing parameters, we found higher rates of editing by increasing donor and a1/a2 length, and using a centered or slightly asymmetric donor with more homology on the PAM-proximal side of the non-target strand. We also demonstrated significant flexibility in the 3’ side of the *msd* sequence for editing, which we altered with targeted deletions, single-nucleotide changes, and stem length alterations. We also changed the 3’ side of the *msd* region to machine learning predicted *de novo* variants of 55-80% difference from the wild-type sequence in the 20 nucleotide ML variable region, or up to 12% over the full retron-Eco1 ncRNA.

Our variant libraries agree with previous optimizations with single-stranded oligonucleotides (ssODNs) in some aspects, and disagree in others. For example, previous work on ssODNs has found that ssODNs of 70-80 nucleotides have the highest rate of precise repair, and precise repair rates decline above 80 nucleotides^24,25^. This is contrary to our finding that precise editing rates increase with increasing length of the RT-DNA past the previously-found optimal length of ssODNs. This difference could be due to lower DNA transfection of longer oligonucleotides or due to the difficulty of synthesizing longer oligonucleotides^24^. As our donor is created inside the nucleus of the cell by the retron-Eco1 RT, our precise editing method will not be limited by synthesis or transfection limitations. We note that, eventually, the retron-Eco1 RT processivity may hinder production of a longer donor, but that we do not believe was have reached that limit in this work, or that any processivity losses are offset by precise repair gains.

Prior optimization work of ssODN donors has also found that donors asymmetric about the cut site on the non-target strand have better precise editing outcomes, agreeing with our results^7,25,26^. After cleavage, Cas9 releases the non-target strand, after which a 3’-to-5’ exonuclease, like Klenow, degrades the 3’ flap^27^. Therefore, homology should be biased and asymmetric towards the PAM-proximal side of the non-target strand, as this strand is both free and non-degraded.

We only evaluated asymmetry in a donor homologous to the non-target strand in this study. This is because, in both yeast and human, across different PAMs, we find donors homologous to the non-target strand result in higher precise editing than the target strand, as fits with the mechanism of Cas9 above and to ssODN studies^24^. However, because our editor is a ncRNA reverse-transcribed into a donor, we have the additional complexity of RNA:RNA hybridization. When the reverse-transcribed donor is homologous to the target strand, the gRNA would be homologous to the donor before reverse transcription and cause the gRNA to be “hidden” from Cas9 through base pairing with the ncRNA donor. This is an additional complexity not evaluated in optimizing ssODNs, and may increase the effect we observe, with non-target strand complementarity of the donor performing better than target strand complementarity.

To our knowledge, this is the first demonstration of using variant libraries to train a ML library that we can query with *de novo* retron ncRNA sequences to assess their possible RT-DNA production. This high-throughput computational approach allowed us to screen many more sequences in silico than currently possible experimentally. Through this, we queried and validated new aspects of the ncRNA that can increase RT-DNA production, and thus editing. Importantly, we were able to use the output of the ML model to make semi-synthetic ncRNAs that are as functional as wild-type.

## Data Availability

All data supporting the findings of this study are available within the article and its supplementary information, or will be made available from the authors upon request. Sequencing data associated with this study are available in the NCBI SRA (PRNJNA1121319). A link for reviewers to access the SRA can be found here: https://dataview.ncbi.nlm.nih.gov/object/PRJNA1121319?reviewer=2b83j6m1ma4osb45ntu4pa070i

## Code Availability

Custom code to process or analyze data from this study will be made available on GitHub prior to peer-reviewed publication: https://github.com/Shipman-Lab/retron_ncRNA_ML_libraries/tree/master

## Acknowledgements

Work was supported by funding from the National Science Foundation (MCB 2137692), the National Institute of Biomedical Imaging and Bioengineering (R21EB031393), and the W.M. Keck Foundation. S.L.S. is a Chan Zuckerberg Biohub - San Francisco Investigator and acknowledges additional funding support from the Pew Biomedical Scholars Program. K.D.C. is supported by a National Science Foundation Graduate Research Fellowship and a UCSF Discovery Fellowship. We would also like to acknowledge the Gladstone Flow Cytometry Core, which performed the fluorescence-activated cell sorting of the human cells in Figure 4.

## Author Contributions

K.D.C. and S.L.S. conceived the study and, with the help of S.C.L., outlined the scope of the project and designed experiments. Experiments were performed and analyzed by S.L.S. (Fig 1, Supp Fig 1), S.L.S. and H.G. (Fig 2), K.D.C. (Fig 3, Supp Fig 2-8), and A.G.K. and K.D.C. (Fig 4). K.D.C. and S.L.S. wrote the manuscript, with input from all authors.

## Competing Interests

L. S. is a co-founder of Retronix Bio and Sprint synthesis.

## Supplementary Figures

**Supplementary Figure 1.**
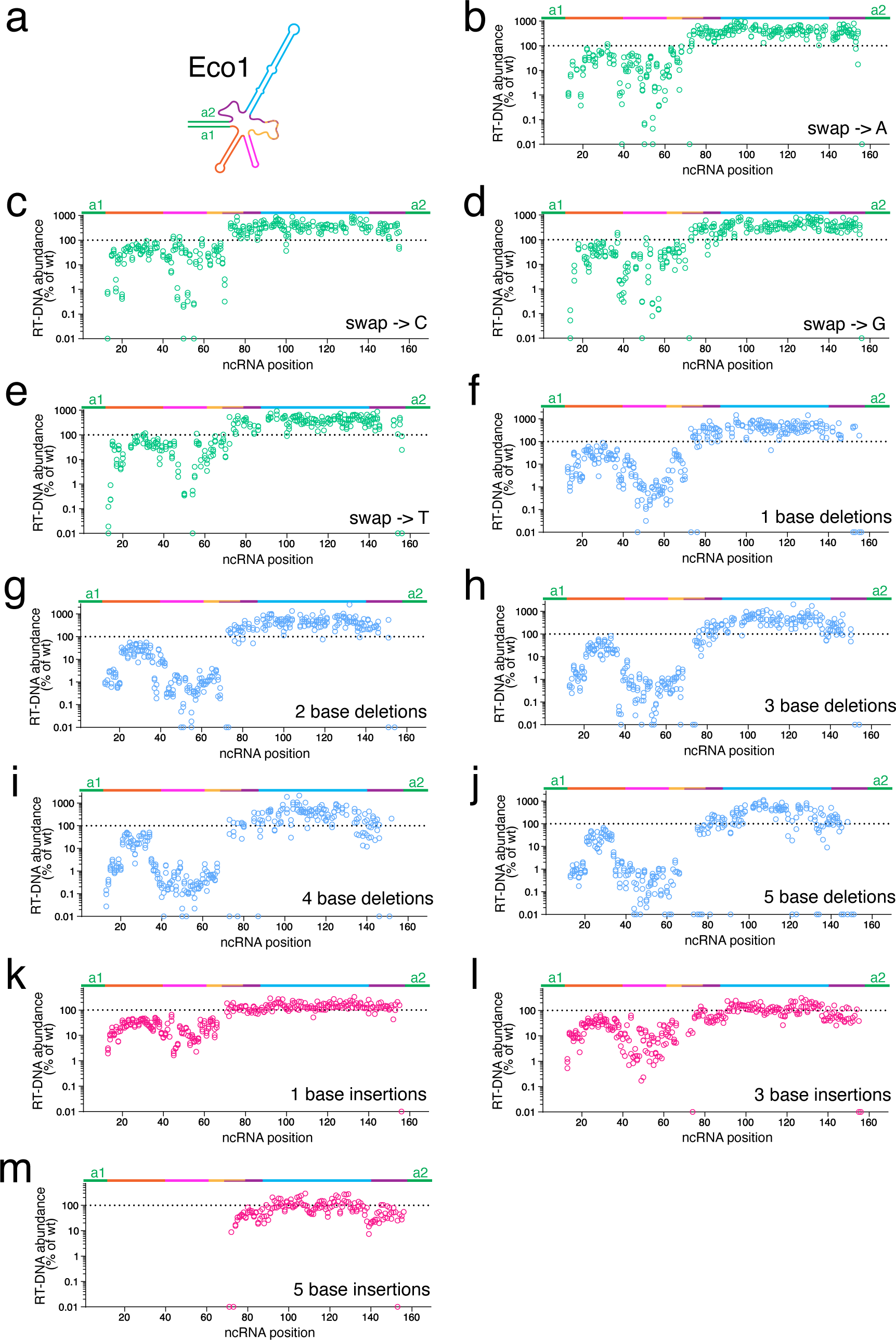
Substitution, deletion, and insertion sub-library RT-DNA production in *E. coli*. **a.** Retron-Eco1 ncRNA structure. **b.** RT-DNA production of N→A nucleotide swap, starting at a specified ncRNA position relative to wild-type RT-DNA. Each open circle represents an individual biological replicate. **c.** RT-DNA production of N→C nucleotide swap, starting at a specified ncRNA position relative to wild-type RT-DNA. Each open circle represents an individual biological replicate. **d.** RT-DNA production of N→G nucleotide swap, starting at a specified ncRNA position relative to wild-type RT-DNA. Each open circle represents an individual biological replicate. **e.** RT-DNA production of N→T nucleotide swap, starting at a specified ncRNA position relative to wild-type RT-DNA. Each open circle represents an individual biological replicate. **f.** RT-DNA production of single-base deletions, starting at a specified ncRNA position relative to wild-type RT-DNA. Each open circle represents an individual biological replicate. **g.** RT-DNA production of two-base deletions, starting at a specified ncRNA position relative to wild-type RT-DNA. Each open circle represents an individual biological replicate. **h.** RT-DNA production of 3-base deletions, starting at a specified ncRNA position relative to wild-type RT-DNA. Each open circle represents an individual biological replicate. **i.** RT-DNA production of 4-base deletions, starting at a specified ncRNA position relative to wild-type RT-DNA. Each open circle represents an individual biological replicate. **j.** RT-DNA production of 5-base deletions, starting at a specified ncRNA position relative to wild-type RT-DNA. Each open circle represents an individual biological replicate. **k.** RT-DNA production of single-base insertions, starting at a specified ncRNA position relative to wild-type RT-DNA. Each open circle represents an individual biological replicate. **l.** RT-DNA production of 3-base insertions, starting at a specified ncRNA position relative to wild-type RT-DNA. Each open circle represents an individual biological replicate. **m.** RT-DNA production of 5-base insertions, starting at a specified ncRNA position relative to wild-type RT-DNA. Each open circle represents an individual biological replicate.

**Supplementary Figure 2.**
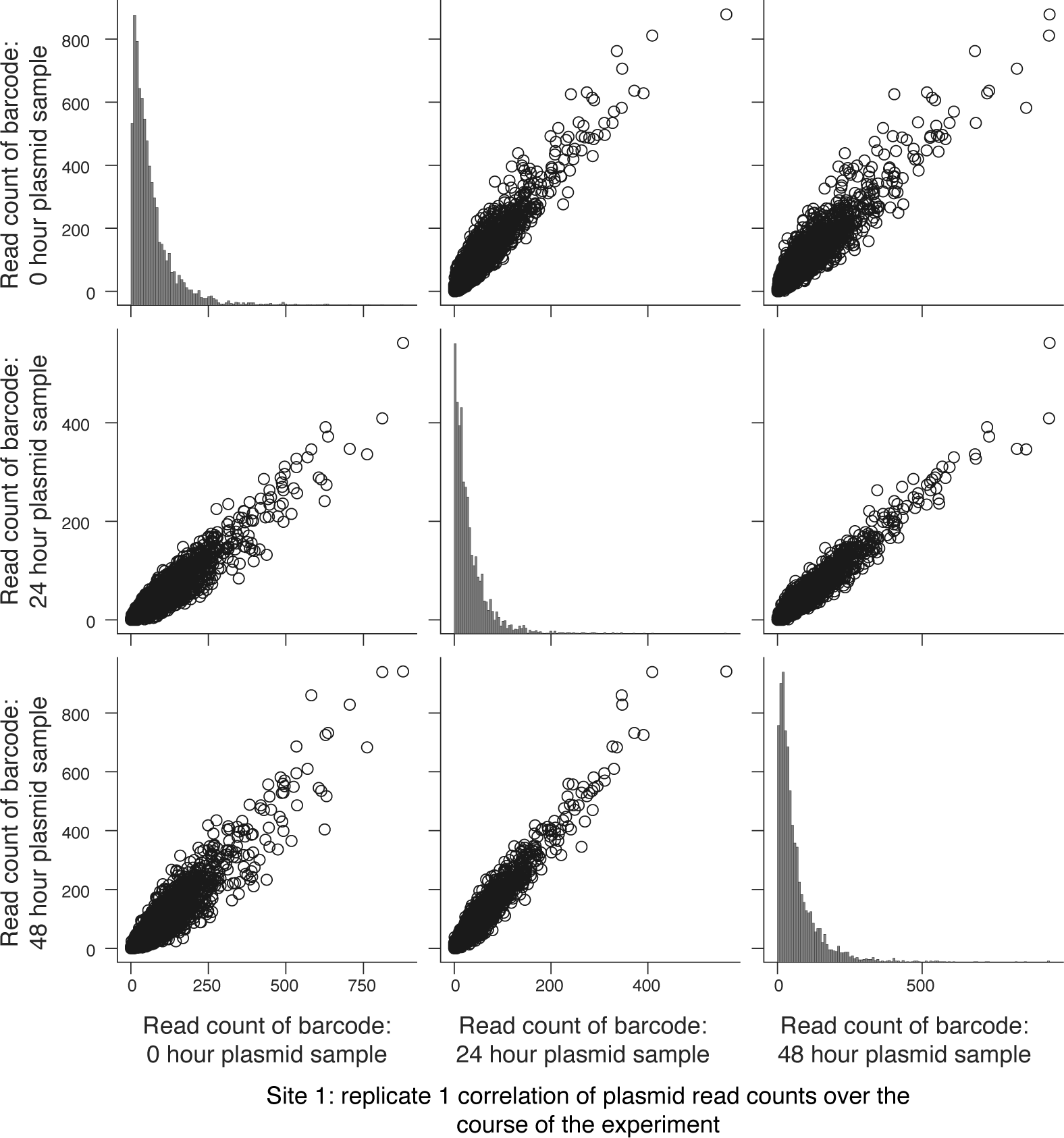
Correlation in plasmid read counts over an example 48-hr editing window in *S. cerevisiae*. Correlation between individual plasmid barcode read counts at 0 hr, 24 hr, and 48 hr of editing for the first biological replicate of the site 1 library. Each open circle represents an individual barcode read count.

**Supplementary Figure 3.**
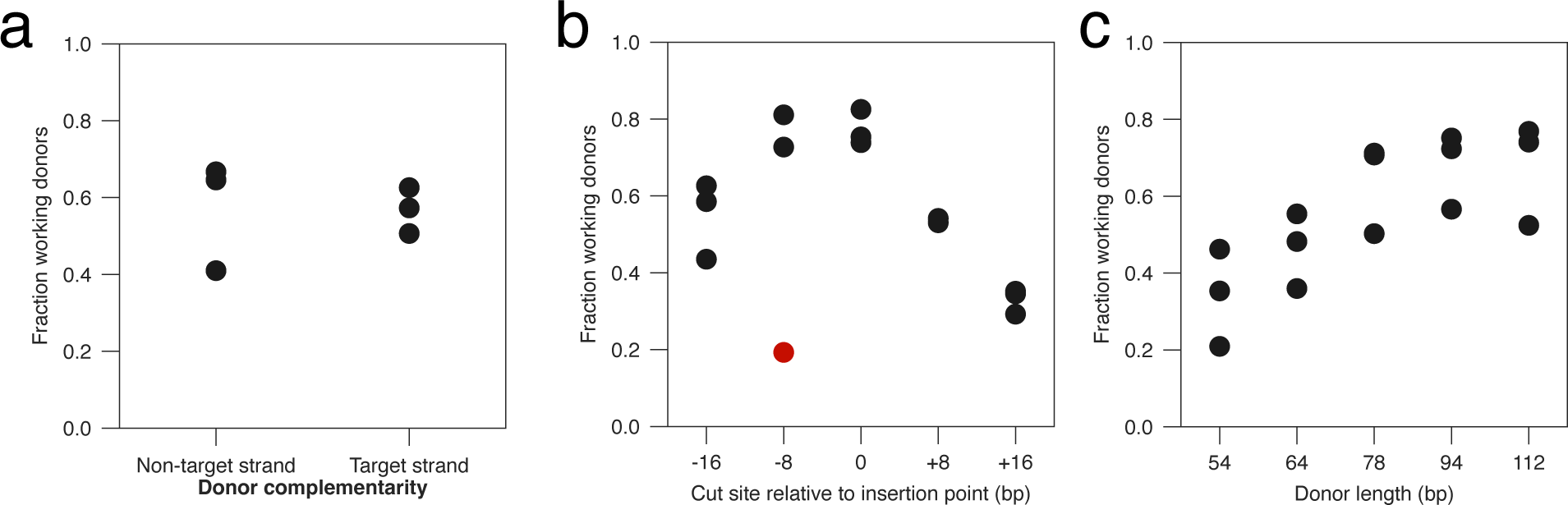
Fraction working editors for different editing variables. **a.** Fraction working donors that are complementary to the non-target strand vs. target strand when reverse-transcribed into donor DNA. Each closed circle represents the mean of the three biological replicates for that site. **b.** Fraction working donors across all tested PAMs. Each closed circle represents the mean of the three biological replicates for that site. The red closed circle represents the PAM excluded from the analysis, as the fraction working donors is below 20%. **c.** Fraction working donors across all tested donor lengths. Each closed circle represents the mean of the three biological replicates for that site.

**Supplementary Figure 4.**
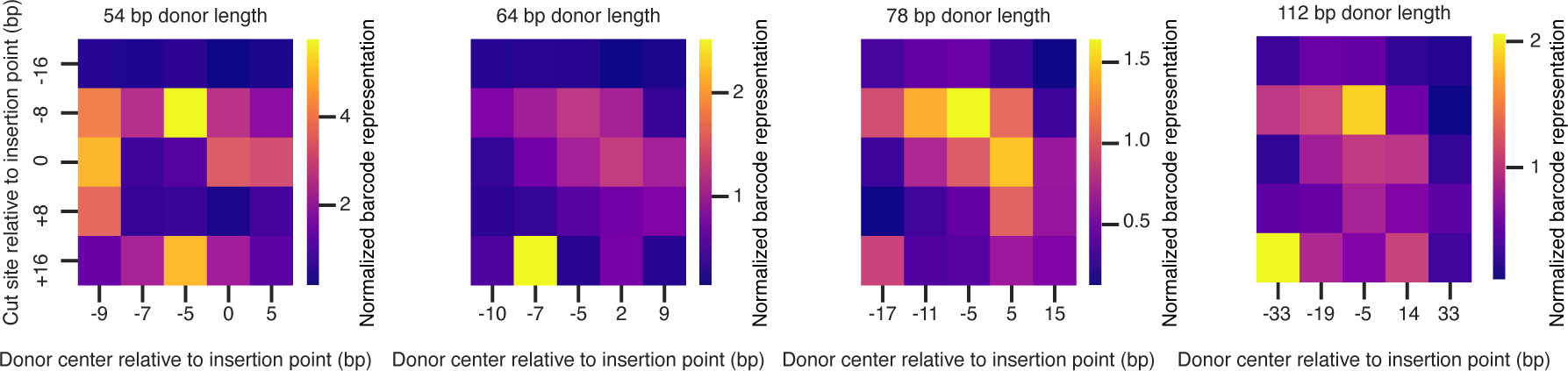
Normalized barcode representation of donors of varying cut sites vs. donor centers in *S. cerevisiae*. Heat map of normalized barcode representation of cut site vs. donor center (54, 64, 78, and 112 nucleotide donor length), normalized to the cut site at the barcode insertion site, and donor center of −5 bp from the barcode insertion site. Each square represents the mean of all biological replicates across all sites.

**Supplementary Figure 5.**
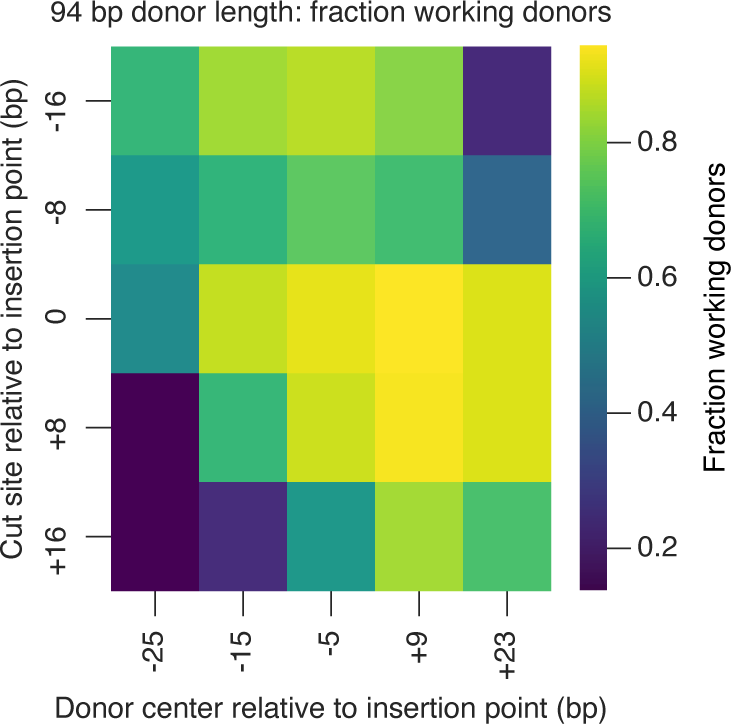
Standard deviation of normalized barcode representation of donors of varying cut sites vs. donor centers in *S. cerevisiae*. Heat map of the standard deviation of normalized barcode representation of cut site vs. donor center (94 nucleotide donor length), normalized to the cut site at the barcode insertion site, and donor center of −5 bp from the barcode insertion site. Each square represents the standard deviation of all biological replicates across all sites.

**Supplementary Figure 6.**
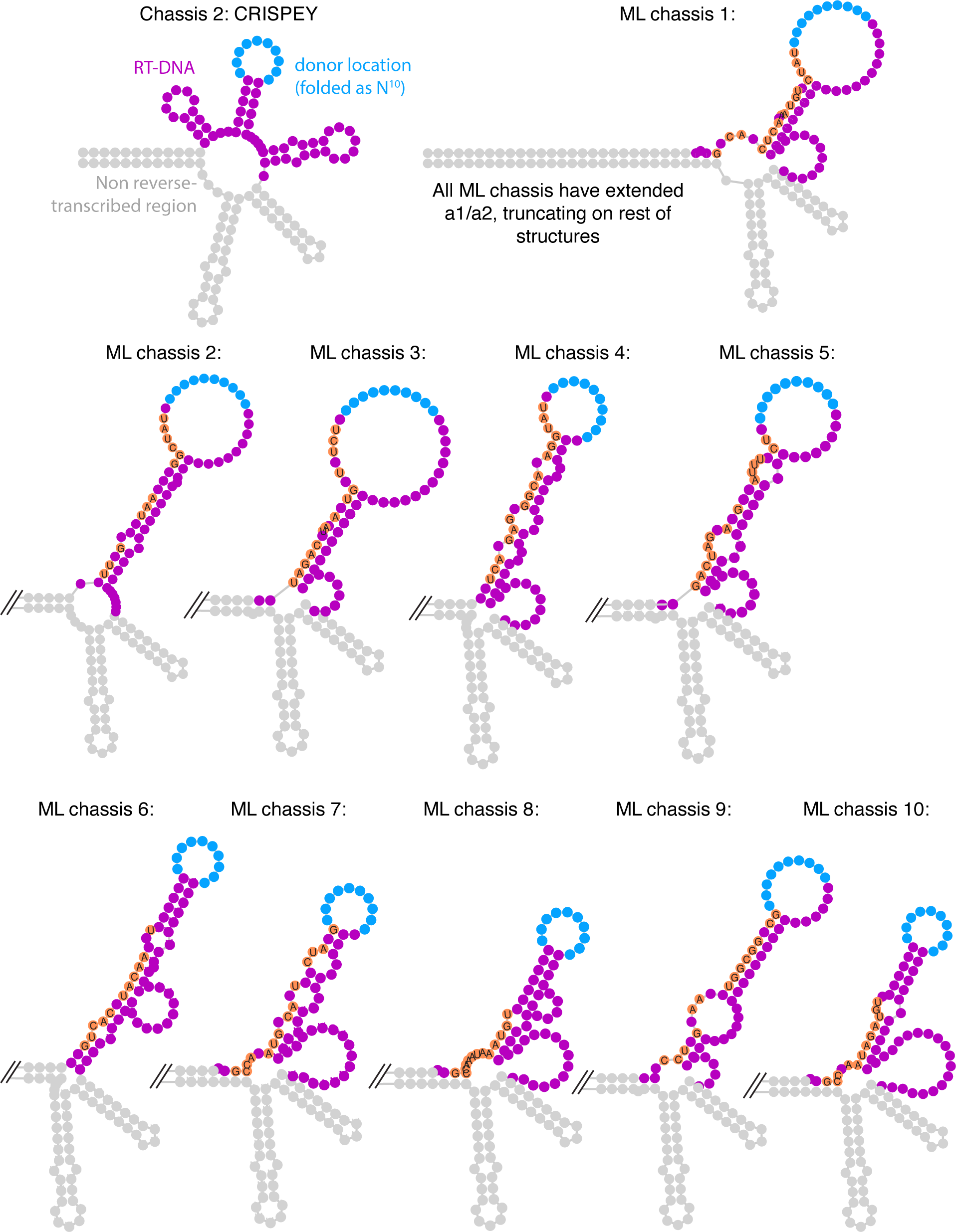
ncRNA structure and sequence of top machine learning ncRNA chassis. CRISPEY and ML chassis were folded using RNAfold (Institute for Theoretical Chemistry, University of Vienna webtool) using N_10_ to stand in for the variable donor region (light blue). *msr* annotated in grey and RT-DNA annotated in purple. Nucleotides with changes from the CRISPEY reference are highlight in red with the nucleotide identity annotated in black.

**Supplementary Figure 7.**
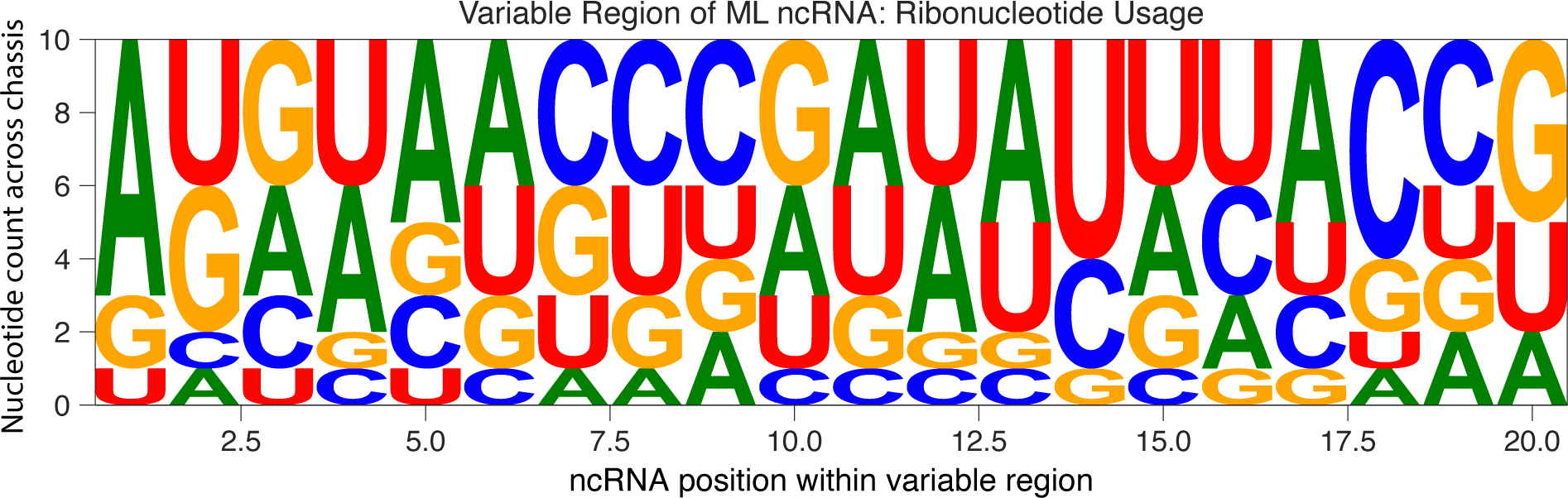
Usage of ribonucleotides in ML ncRNA chassis across variable region.

**Supplementary Figure 8.**
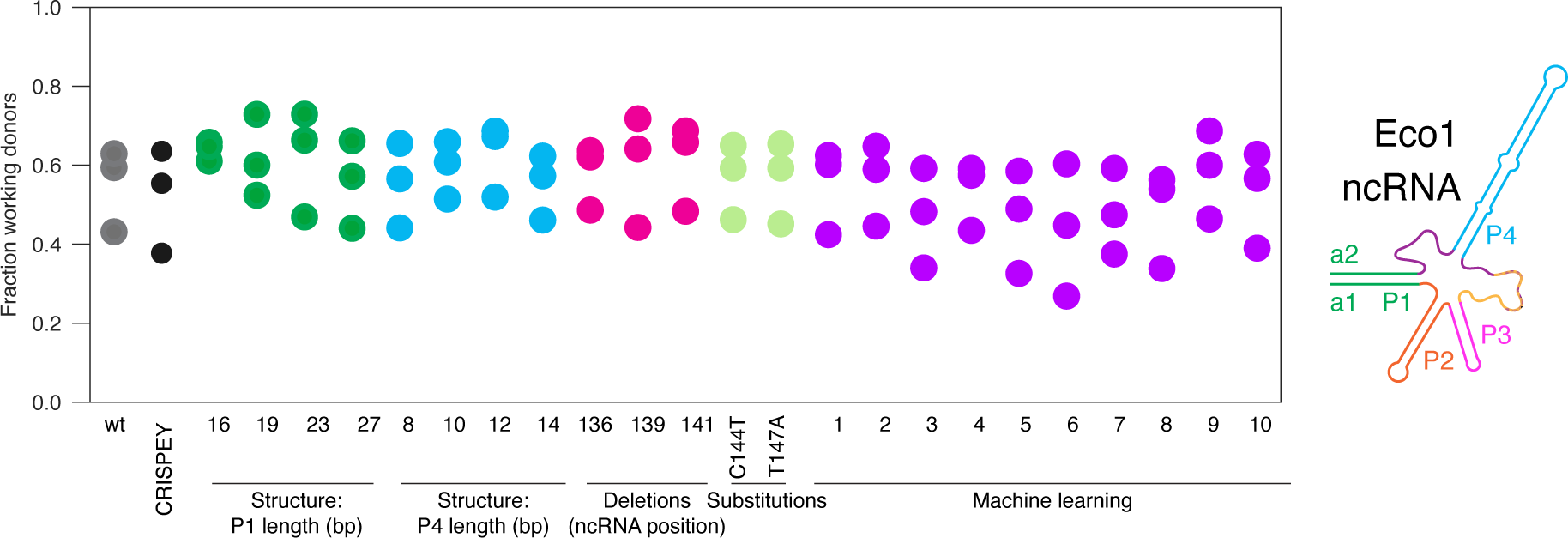
Fraction working editors across ncRNA chassis. Fraction of working donors across all ncRNA chassis. Each closed circle represents the mean of the three biological replicates for that site.

## Notes

https://github.com/Shipman-Lab/retron_ncRNA_ML_libraries/tree/master

